# An interaction-based model for neuropsychiatric features of copy-number variants

**DOI:** 10.1101/459958

**Authors:** Matthew Jensen, Santhosh Girirajan

## Abstract

Variably expressive copy-number variants (CNVs) are characterized by extensive phenotypic heterogeneity of neuropsychiatric phenotypes. Approaches to identify single causative genes for these phenotypes within each CNV have not been successful. Here, we posit using multiple lines of evidence, including pathogenicity metrics, functional assays of model organisms, and gene expression data, that multiple genes within each CNV region are likely responsible for the observed phenotypes. We propose that candidate genes within each region likely interact with each other through shared pathways to modulate the individual gene phenotypes, emphasizing the genetic complexity of CNV-associated neuropsychiatric features.

## A case for a multi-genic model of CNV pathogenicity

Since the advent of large-scale sequencing studies, the number of genes associated with neurodevelopmental disorders such as autism, intellectual disability, and schizophrenia has increased dramatically. For example, nearly 200 genes have been identified with recurrent *de novo* mutations in both individuals with autism and intellectual disability (1–8). In fact, complex human disease phenotypes can be influenced by variation in both a small number of core genes with large effect size and a large number of modifier genes with small effect size, accounting for the large number of candidate neurodevelopmental genes (9,10). The application of a multi-genic model for disease pathogenicity has not been fully expanded to cover copy-number variants (CNVs), or large duplications and deletions in the genome. The prevailing notion of single causative genes for CNV disorders is due to the paradigm of gene discoveries for CNVs associated with genetic syndromes in individuals with specific constellations of clinical features, such as Smith-Magenis syndrome (SMS). Although some variability in phenotypic expression has been documented, these disorders usually occur *de novo* and are characterized by high penetrance for the observed phenotypes (11,12) (**Figure 1**). In these cases, individuals manifesting the characteristic features of the syndrome but with either atypical breakpoints or mutations in individual genes within the CNV region were used to identify causative genes for the major phenotypes (13–15). These causative genes, such as *RAI1* for SMS, were then confirmed by recapitulating conserved phenotypes of the deletion using functional evaluations in animal models (16,17).

**Figure 1.**
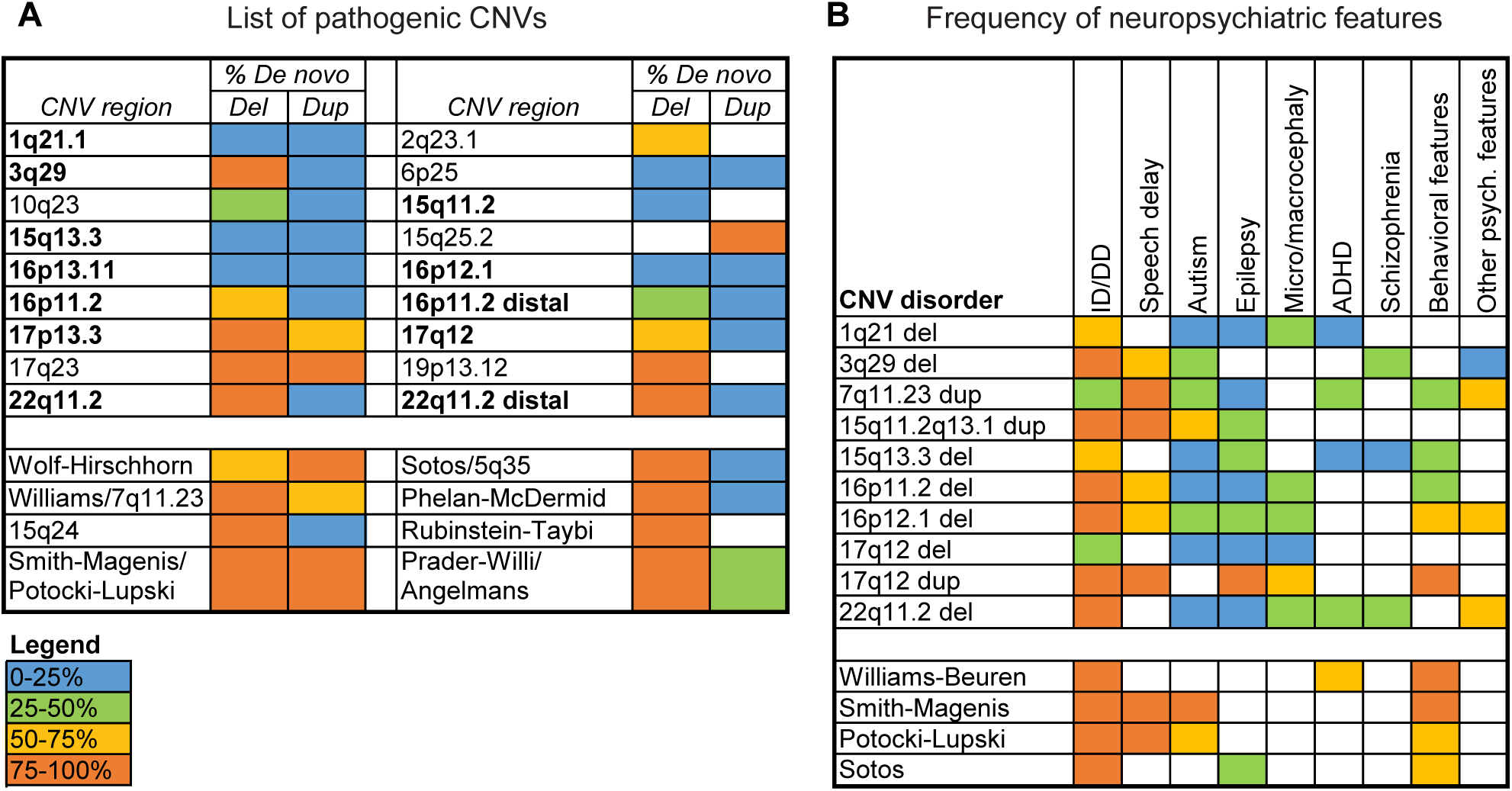
Phenotypic profiles of syndromic and variably expressive CNVs. **(A)** Table listing variably expressive (top) and syndromic (bottom) CNV regions is shown. The colored boxes indicate frequency of *de novo* versus inherited CNV cases for deletions (del) and duplications (dup) previously identified in a cohort of 2,312 children with developmental disorders (12). The twelve variably-expressive CNV regions highlighted in bold were selected for the analysis described in the manuscript. **(B)** Table listing average frequencies of neurodevelopmental phenotypes for select variably-expressive and syndromic CNVs, curated from GeneReviews reports on individual CNVs (107), is shown. White boxes represent no available data from GeneReviews, but do not necessarily indicate a lack of association between the CNV and the phenotype (for example, 1q21.1 deletion and schizophrenia). Data for this figure are available in the **Supporting Information** file.

In contrast, another category of CNVs has been identified in individuals with neurodevelopmental disorders, including duplications and deletions at proximal 16p11.2, 3q29, distal 16p11.2, and 1q21.1 (18–21). Although these CNVs are enriched in affected individuals compared to population controls, they are primarily characterized by variable expressivity of clinical features (12,22–26) (**Figure 1B**). For example, the 16p11.2 deletion has been implicated in 1% of individuals with idiopathic autism (18,27), but only 25% of individuals with the deletion exhibit an autism phenotype (28–31), while others may manifest intellectual disability, obesity, or epilepsy at varying degrees of penetrance (28,32,33). In fact, certain CNVs, such as the 16p12.1 deletion and the 15q11.2 deletion, have a high frequency of carriers who only manifest mild neuropsychiatric features, in contrast to more severely affected individuals who also carry other rare variants in the genetic background (12,22,23,26,34,35). As such, many variably expressive CNVs have a higher frequency of inherited compared to *de novo* occurrence (12) (**Figure 1A**).

Based on the success of gene discovery in CNVs with syndromic features, such as SMS, several studies have attempted to identify the causative genes in variably expressive CNVs (36–52). Several individual genes within variably expressive CNV regions have been associated with specific congenital or structural features of these disorders, including *TBX6* for scoliosis in 16p11.2 deletion (53), *TBX1* for cardiac phenotypes in 22q11.2 deletion (38,54), *GJA8* for cataracts and *GJA5* for heart defects in 1q21.1 deletion (55,56), and *MYH11* for aortic aneurysms in 16p13.11 duplication (57,58). However, approaches to identify single causative genes for the more prominent neuropsychiatric features of these CNVs have not been successful (59). Here, we show several lines of evidence from gene pathogenicity metrics, animal model studies, and gene expression data that support the involvement of multiple genes towards the neuropsychiatric features of variably expressive CNVs.

*First*, genome-wide metrics of pathogenicity, including those that measure haploinsufficiency (HI score, gene essentiality, GHIS and EpiScore) (60–63) and resistance to variation (RVIS, pLI and maximum CCR scores) (64–66), provide evidence for several candidate genes within CNV regions for developmental disorders **(Figure 2)**. For example, 45 out of 152 genes (30%) within 12 variably expressive CNV regions are intolerant to variation with RVIS metrics in the top 20^th^ genome-wide percentile, similar to that of known neurodevelopmental genes such as *CHD8, NRXN1* and *SCN2A*, as well as genes responsible for major features of syndromic CNVs, such as *RAI1* and *NSD1* (**Figure 2A**). These top-ranked genes include *TAOK2*, *MVP, ALDOA* and *DOC2A* on chromosome 16p11.2, *BCL9* and *GJA5* on chromosome 1q21.1, and *ATXN2L*, *ATP2A1* and *SH2B1* on distal 16p11.2. Similarly, 32/165 genes (19%) are considered intolerant to loss-of-function mutations based on pLI scores (>0.9), and 36/160 genes (23%) have haploinsufficiency scores in the highest 20^th^ percentile of the entire genome (**Figure 2A**). Further, the top 10% of all genes identified by a gene interaction-based machine learning classifier to be associated with autism included eight genes within 16p11.2 and four genes within 22q11.2 (67).

**Figure 2.**
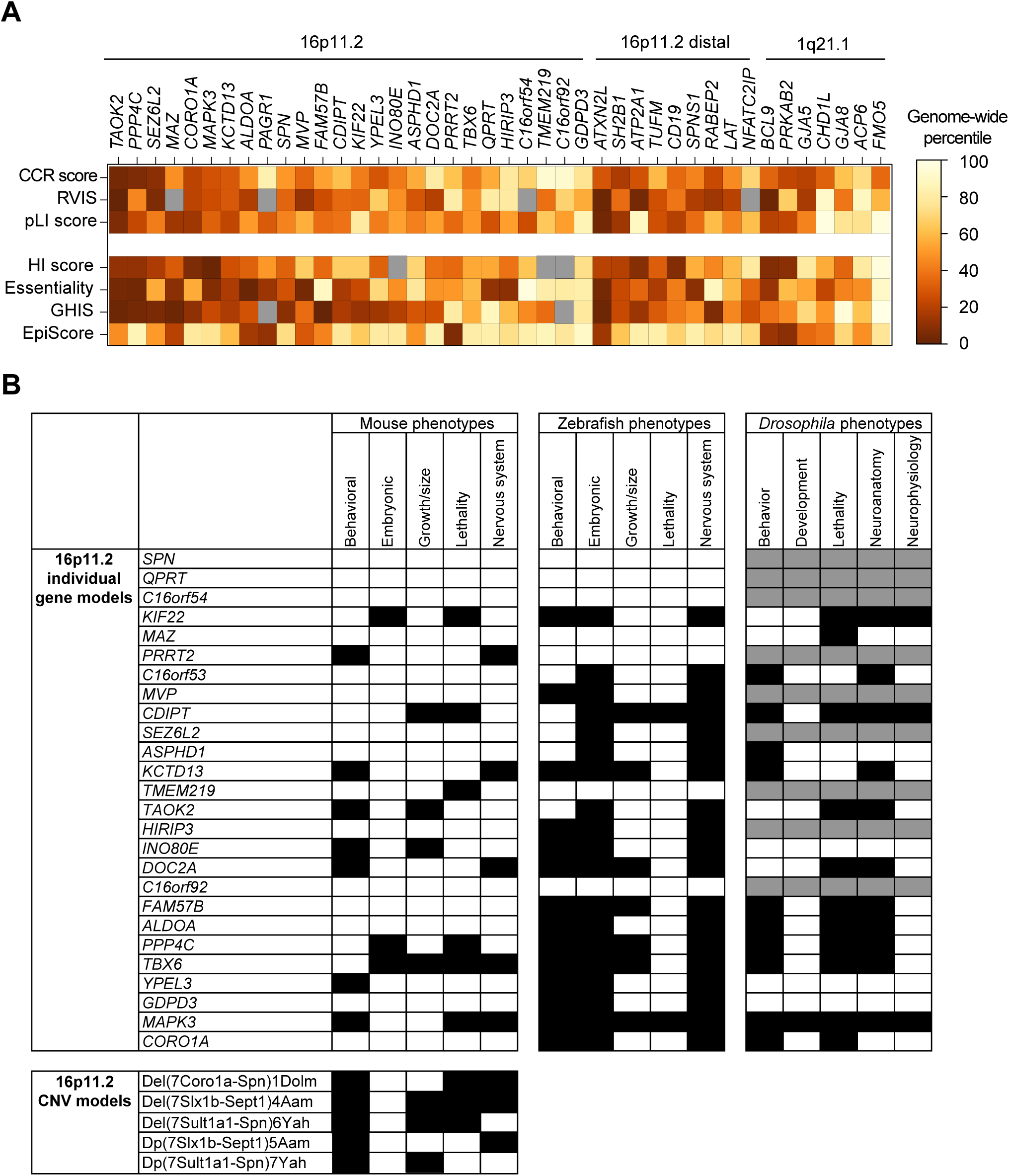
**(A)** Percentile-rank scores compared to the whole genome for intolerance to variation (RVIS, pLI and maximum CCR) and haploinsufficiency (HI, essentiality, GHIS and EpiScore) metrics for genes within select variably expressive CNV regions (60–66). Lower percentile scores indicate a gene is more likely to be haploinsufficient or intolerant to variation. Grey boxes indicate metrics were not available for a particular gene. **(B)** Developmental phenotypes in animal models for homologs of individual genes within the 16p11.2 region, as catalogued from animal model databases (MGI, ZFIN and FlyBase). Black boxes indicate presence of phenotype, white boxes indicate absence of phenotype, and grey boxes indicate no homolog is present for a particular gene in a model organism. The phenotypes observed in 16p11.2 deletion and duplication mice are distinct from those observed in the individual gene models (81–85). Data for this figure, including gene metrics and animal phenotypes for other CNV genes not shown in this figure, are available in the **Supporting Information** file. (Abbreviations: RVIS—Residual Variance to Intolerance Score; pLI—Probability of Loss-of-function Intolerance; CCR— Constrained Coding Regions; HI—Haploinsufficiency score, GHIS—Genome-wide haploinsufficiency score; MGI—Mouse Genome Informatics; ZFIN—Zebrafish Information Network)

*Second*, several recent studies using animal and cellular models have demonstrated the critical involvement of several genes within CNVs towards neurological, cellular and developmental functions (36,37,46,47,51,52) (**Figure 2B**). For example, Blaker-Lee *et al*. screened 22 homologs of 16p11.2 genes in zebrafish morpholino knockdown models, and identified 20 homologs that contributed to morphological defects and abnormal behavior (37). Iyer *et al.* also screened homologs of 16p11.2 genes in *Drosophila melaogaster* using RNAi knockdown, and found that 10 out of 14 homologs contributed to global developmental defects as well as specific neuronal and cellular defects in the developing fly eye (46). Further, mouse models for 15 genes within the 16p11.2 region have been generated to test for defects in development and neuronal behavior (45,48–50,68–80). For example, *Taok2*^-/-^ mice have increased brain size, behavioral defects, and impaired synapse development (50), *Kcdt13*^+/-^ mice show defects in hippocampal synaptic transmission and decreased dendritic complexity (45), *Mapk3*^+/-^ mice show behavior anomalies, abnormal synapse function and reduced cell proliferation during development (68,69), and *Mvp*^+/-^ mice show decreased plasticity and synaptic defects in ocular neurons (48) (**Figure 2B**). Importantly, these models of individual genes do not fully recapitulate the phenotypes observed in models of the entire CNV (81–85). For example, the decreased body weight, abnormal brain morphology and coordination defects observed in 16p11.2 deletion mouse models have not been observed in any individual gene knockdown models (81–84) (**Figure 2B**). Similarly, *Otud7a^+/-^* mouse models have low body weight, reduced vocalization, abnormal dendritic spine morphology, and seizures, but the 15q13.3 deletion mice also show learning and memory defects in addition to the above features (43,44,86). Further, mouse models for *Chrna7^+/-^*, another candidate gene on chromosome 15q13.3, only show subtle behavioral phenotypes (87). These data suggest that haploinsufficiency of *CHRNA7* or *OTUD7A* alone is not sufficient to account for the pathogenicity of the entire CNV. Overall, a catalog of functional data from mouse (88), zebrafish (89), and fruit fly studies (90) indicates that 80% (131/163) of homologs for genes within CNV regions present lethality, behavioral, developmental, or neuronal phenotypes when disrupted. These data suggest that disruption of multiple genes within each CNV region can affect important developmental or neuronal functions that could contribute to the phenotypes of the entire CNV.

*Third*, patterns of gene expression in humans and model organisms have identified multiple genes within each CNV region that are co-expressed in the developing brain along with known neurodevelopmental genes. For example, Maynard and colleagues examined expression patterns of 22q11.2 gene homologs in the developing mouse brain, and found that 27 out of 32 genes were expressed in the embryonic forebrain, with six genes expressed in neuronal tissues related to schizophrenia (39). In fact, a genome-wide weighted gene correlation network analysis (WGCNA) (91) from different brain tissues during development (92) shows several large modules of genes with similar expression patterns (**Figure 3**). For example, the five largest modules are each enriched (p<0.05 with Benjamini-Hochberg correction) for biological functions related to neurodevelopment, including protein modification and transport in module 1 (M1), nervous system development in M2, and cell communication and signal transduction in M5. Importantly, each of these modules contains multiple genes from the same CNV region, including 3q29 genes *PAK2*, *NCBP2*, and *BDH1* in M1, 1q21.1 genes *BCL9, CHD1L* and *FMO5* in M2, and 16p11.2 genes *MVP* and *QPRT* in M5. Therefore, it is clear that multiple genes in the same CNV region are co-expressed with each other in the developing brain and could share similar functions or regulatory patterns.

**Figure 3.**
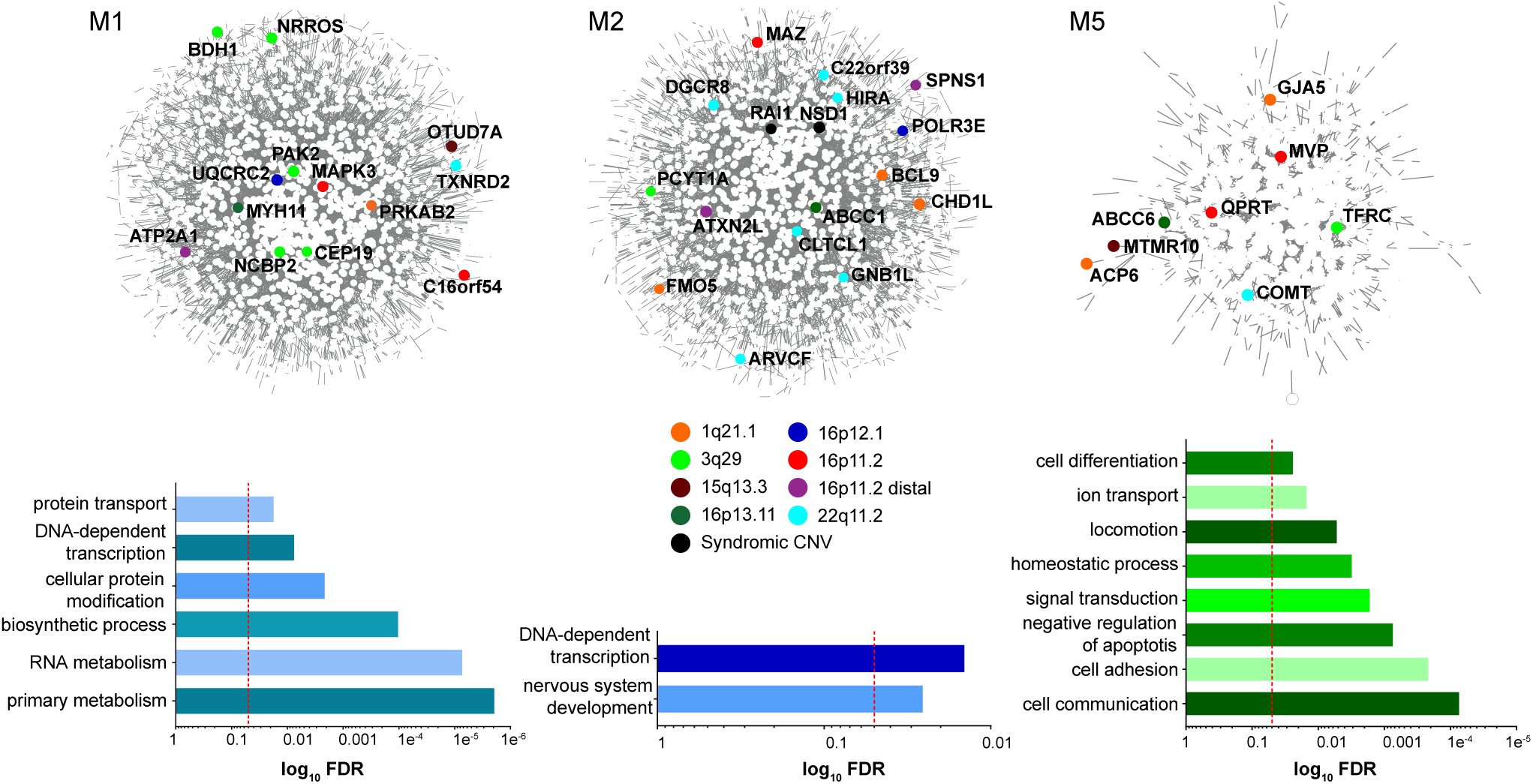
Modules of co-expressed genes derived from WGCNA analysis of BrainSpan Atlas RNA-Seq data (Gencode v10) (92) across 524 tissues and timepoints the developing brain. Networks of interactions among genes within three select top WGCNA modules (M1, M2 and M5) were obtained from the BioGrid interaction database (108) and visualized with Cytoscape (109). Genes within variably expressive CNV regions are highlighted as colored nodes in each network. Bar graphs show enrichment (p<0.05 with Benjamini-Hochsberg correction, represented by red dotted line) of genes within each module for Gene Ontology (GO) Biological Process terms, calculated using PantherDB (110). Data for this figure are available in the **Supporting Information** file.

## Dissecting the genetic complexity of CNV pathogenicity

Several scenarios could explain how the haploinsufficiency of multiple genes can predict the variable phenotypes associated with the entire CNV (**Figure 4A**). The simplest such model is an additive model, where disruption of individual genes within a CNV may only impart a mild phenotype on their own, but additively contribute to more severe features (93) (**Figure 4A**). However, an additive model may not always explain the phenotypic features manifested by CNVs containing multiple candidate genes that could lead to severe defects or lethality on their own. For example, heterozygous *Tbx1^+/-^* (within the 22q11.2 region) and *Mapk1*^+/−^ (within the distal 22q11.2 region) mice both lead to perinatal or neonatal lethality (94–96). In humans, 14% (24/172) of CNV genes are under evolutionary constraint in control populations (pLI score >0.9 or maximum CCR score >99^th^ percentile) and have no reported disease-associated variants (97–99), suggesting that these genes could be under strong purifying selection (66). Further, 18% (22/125) of CNV genes show evolutionary constraint for loss-of-function mutations (pLI>0.9) but not for copy-number changes within a control population (100). We therefore hypothesize that the pathogenicity of variably expressive CNVs can also be explained by complex interactions among the constituent genes within shared biological pathways. These interactions can enhance or suppress the phenotypes caused by disruption of individual genes. Under this model, the haploinsufficiency of certain genes can be modulated by haploinsufficiency of other interacting genes in the same region that may or may not lead to phenotypes on their own (**Figure 4A**). Further, variants in the genetic background that map within these shared pathways can simultaneously modulate the effects of multiple genes, ultimately defining the phenotypic trajectory in CNV carriers (**Figure 4A**). For example, Pizzo *et al.* found that the burden of rare deleterious mutations within genes in the genetic background correlated with variability of IQ scores and head circumference among 16p11.2 deletion carriers (35). The potential for complex interactions within a CNV region depends on the functional convergence of the constituent genes. For instance, both *KCTD13* and *TAOK2* within 16p11.2 participate in the RhoA signaling pathway (45,50) and therefore are more likely to interact with each other than genes located in different biological pathways. In fact, it has been shown that genes within pathogenic CNVs are more similar in function compared to genes within benign CNVs, suggesting that variably expressive CNVs are likely to contain interactions between functionally relevant genes (101). Further, Noh and colleagues found an over-representation of interactions among genes within autism-associated CNVs, and these interactions were enriched for synaptic transmission and regulatory signaling pathways (102). Because of this, therapeutic targets for pathways shared among CNV genes could be explored as potential treatments for CNV disorders.

**Figure 4.**
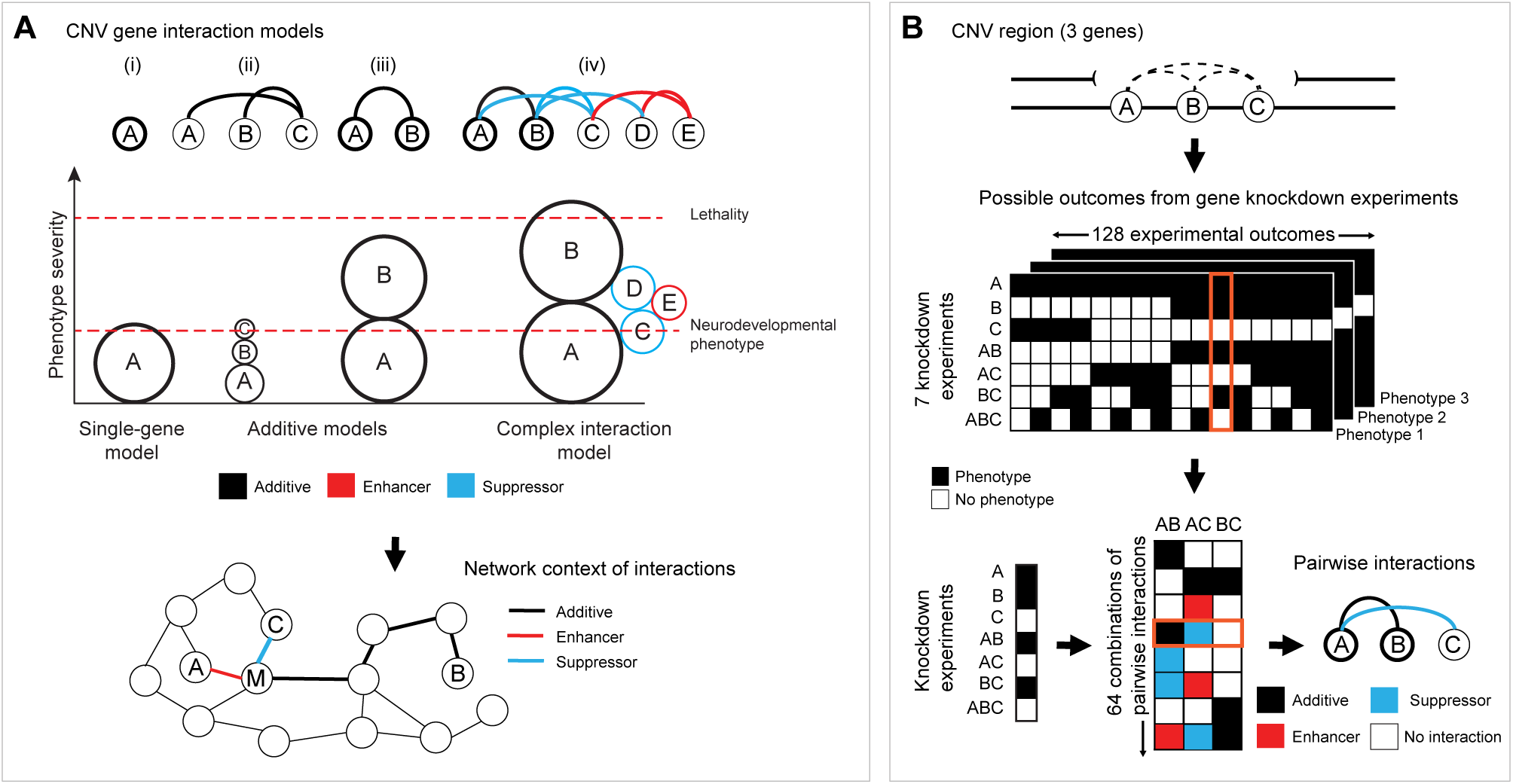
Models for genetic interactions within CNV regions. **(A)** Several models of interactions among CNV genes are shown. These models include (i) a single-gene model where one gene is sufficient to account for the phenotype; additive models where the phenotype is due to the additive effects of multiple CNV genes that (ii) may or (iii) may not account for phenotypes on their own; and (iv) a complex interaction model where additive, enhancer and suppressor interactions between genes in the CNV region modulate the phenotype, including when additive effects could lead to lethality. The size of the circles in the plot indicates the relative contribution of each gene to the overall neurodevelopmental phenotype. Thick circles indicate genes that contribute to the observed phenotypes on their own, while connector lines indicate the nature of interaction between pairs of genes. Connected modifier genes (M) can further modulate these interactions to ultimately define the phenotypic trajectory in individuals carrying the CNV. **(B)** For a hypothetical CNV region with three genes, there are seven combinations of gene knockdowns (A, B, C, AB, BC and ABC) that can be tested for the presence or absence of a specific phenotype. These knockdown experiments can yield 128 potential outcomes for each phenotype tested, with each individual set of outcomes corresponding to one of 64 combinations of pairwise gene interactions (additive, enhancer, suppressor or no interaction). One possible outcome highlighted in orange shows presence of a particular phenotype for knockdowns of single genes A and B and two-hit knockdowns AB and BC. The single-gene knockdowns indicate that only genes A and B contribute to the phenotype, and that the phenotype of pairwise knockdown AB is due to the additive effects of the two genes. While the phenotype is observed for BC, the phenotype is not observed for AC and ABC, suggesting that gene C suppresses the phenotype of gene A.

The possibility of additive, suppressor and enhancer interactions between pairs of genes underlies the potential for highly complex models of CNV pathogenicity. For instance, within a CNV region spanning three genes, seven combinations of gene knockdown experiments (haploinsufficiency of A, B, C, AB, BC, AC, and ABC) can be tested for the presence or absence of a specific phenotype (**Figure 4B**). This set of knockdown experiments can yield 128 possible experimental outcomes that can be used to further deduce 64 possible sets of pairwise interactions for AB, BC, and AC (no interaction, additive, suppression, or enhancement for each interaction) (**Figure 4B**). These possible combinations of interactions exponentially increase for larger CNVs with more genes, and the complexity further increases if quantitative phenotypes are used to determine the magnitude of interactions between genes or when interactions with variants in the genetic background are taken into account. However, testing even a small number of these interactions would still uncover the nature of the relationships among genes within a CNV region and potentially a common pathway shared by those genes. For example, Grice and colleagues used *D. melanogaster* RNAi models to identify six synergistic interactions out of 41 tested pairwise interactions between genes within *de novo* CNVs from autism patients, including partial 3q29 and 22q11.2 deletions (103). Iyer *et al*. also used fly models to identify 24 additive, enhancer and suppressor interactions out of 52 tested pairwise interactions among homologs of 16p11.2 genes (46), providing further evidence for complex interactions within CNV regions. Further, these interaction models for CNV pathogenicity can be tested in cellular models of the entire CNV. For example, a more severe phenotype observed by restoring dosage of a candidate gene would suggest that disruption of this gene potentially suppresses the effects of other genes within the CNV.

## Complex genetic interactions in the context of genome sequencing

In recent years, exome and whole-genome sequencing analysis has proven invaluable in identifying candidate genes for neurodevelopmental disorders (104). However, sequencing studies would not be able to capture the genetic complexity of a multi-genic CNV region. For example, genes that cause severe phenotypes or lethality on their own and are modulated by haploinsufficiency of other interacting genes within a CNV are less likely to have an enrichment of mutations in sequencing studies. Further, because of the strong phenotypic heterogeneity of these CNVs, it is not possible to determine whether the phenotypes of any individual candidate gene fully recapitulate the variable phenotypes of the entire CNV region. Candidate genes within CNVs identified through genome sequencing studies, such as *TAOK2* on chromosome 16p11.2 (50) or *CHRNA7* on chromosome 15q13.3 (105), do not preclude the possibility of other candidate genes in the same region. Because of this, a thorough systems-based approach for each gene within a CNV and its interactions is necessary to identify candidate genes responsible for the neuropsychiatric features of each region (106).

In summary, genomic and functional data have implicated multiple genes in variably expressive CNV regions towards neuropsychiatric phenotypes, suggesting that single causative genes are not responsible for the heterogeneous features of these CNVs. Here, we propose a complex interaction-based model for these CNVs, where candidate genes within each region interact with each other to influence the variable clinical outcome. The CNV phenotype is therefore distinct from the phenotype manifested by any individual gene, or in some cases, the additive effects of all genes in the region. This multi-genic model of CNVs agrees with a broader complex genetic view of neurodevelopmental disorders, where hundreds of genes with varying effect sizes and complex interactions influence developmental features (10). Further studies on the role of individual genes in CNV regions towards neurodevelopment, especially those that identify key interactions between genes, will be useful in uncovering the cellular pathways and mechanisms responsible for the observed neuropsychiatric features.

## Acknowledgements

The authors thank Lucilla Pizzo, Vijay Kumar and Maitreya Das for their helpful discussions and comments on the manuscript.

## Contributions

M.J. and S.G. conceptualized and wrote the Viewpoint article.

